# Bone Marrow Adipokine Mediates Hematopoietic Regeneration and Stem Cell Fitness

**DOI:** 10.1101/2025.08.28.672647

**Authors:** Parash Prasad, Angelo D’Alessandro, Abhishek K. Singh, Yi Zheng, John Manis, Li Chai, Mohd Kamran, Mark Soliman, Ashley M. Wellendorf, Leo Mejia, Hongbo R. Luo, Sean R Stowell, Shin-Young Park, Martin J. Aryee, Mercedes Ricote, Nathan Salomonis, Jose A. Cancelas

**Affiliations:** Depts. of Medical Oncology, Dana-Farber Cancer Institute, Harvard Medical School, Boston, MA; Department of Biochemistry and Molecular Genetics, University of Colorado-Anschutz, Denver, CO; Division of Experimental Hematology and Cancer Biology, Cincinnati Children’s Hospital Medical Center, Cincinnati, OH; Department of Pediatrics, University of Cincinnati College of Medicine, Cincinnati, OH; Department of Pathology, Boston Children’s Hospital, Harvard Medical School, Boston, MA; Department of Pathology, Mass General Brigham, Harvard Medical School, Boston, MA; Department of Statistics, Dana-Farber Cancer Institute, Harvard Medical School, Boston, MA; Department of Immunology and Oncology, Centro Nacional de Biotecnología (CNB), Consejo Superior de Investigaciones Cientificas (CSIC), Madrid, Spain; Division of Biomedical Informatics, Cincinnati Children’s Hospital Medical Center, Cincinnati, OH

**Keywords:** Hematopoietic stem cells, bone marrow adipocytes, Resistin, NF-kB

## Abstract

Bone marrow (BM) hematopoietic stem cells (HSCs) are exquisitely sensitive to cues from the BM microenvironment (ME), which is critical for their engraftment and regeneration following myeloablative stress. Retinoic acid signaling, acting on both HSCs and niche cells, has emerged as a central regulator of this process. Among ME components, BM adipocytes (BMAs), which can comprise up to 45% of BM volume and expand dramatically during the pancytopenic phase after myeloablation, play a previously underappreciated role in hematopoietic recovery. Here, we identify retinoid X receptor (RXR) signaling in BMAs as a key regulator of the adipokine Resistin, which promotes HSC self-renewal and functional fitness by activating NF-κB signaling. Conditional loss of RXR in adiponectin-expressing cells suppressed Resistin production, resulting in reduced NF-κB activity in HSCs, impaired self-renewal, and defective multilineage hematopoietic regeneration. Functionally, in vivo Resistin neutralization impaired hematopoietic reconstitution, whereas supplementation with either monomeric or dimeric Resistin enhanced HSC self-renewal and long-term lympho-hematopoietic reconstitution in an NF-κB–dependent manner. Together, these findings establish BMA-derived Resistin as an RXR-dependent, critical extrinsic regulator of HSC self-renewal and regenerative hematopoiesis, underscoring its essential role in lympho-myeloid reconstitution after myeloablation.

**Disclosures**: The authors declare no relevant conflicts of interest.

## Introduction

Hematopoietic stem cells (HSC) are a rare population of somatic stem cells whose function is tightly regulated by environmental cues within the bone marrow (BM) niche. Within this niche, HSC receive essential regulatory signals from different microenvironmental cell populations^1,2,3^. BM adipocytes (BMA) represent a major component of the BM cavity, representing up to 45% of the BM volume^4,5^. Like other stromal cell elements, BMAs can influence hematopoiesis through the secretion of soluble factors, yet their overall impact on HSC activity remains controversial, with evidence supporting both inhibitory and supportive roles ^6–9^. Interestingly, the BMA content increases transiently in hematopoietic active BM regions following stressors such as chemotherapy, before returning to baseline levels once regeneration is complete, an observation that suggests that BMAs may provide supportive clues during BM hematopoietic recovery^9,10^.

A key regulator of the BM hematopoietic function during stress is retinoic acid (RA), which signals through nuclear receptors ^11,12^. Retinoid X Receptors (RXRs) mediate RA signaling and function as homodimers or as obligate heterodimer partners with retinoic acid receptors (RARs) and peroxisome proliferator-activated receptor gamma (PPARγ), thereby enabling transcriptional activation of RA-responsive genes^13^. RXR signaling is also indispensable for adipocyte differentiation through its partnership with PPARγ^14^. Based on these insights, we hypothesized that BMA RXR positively regulates hematopoiesis by controlling the expression and/or secretion of adipokines with functional relevance to HSC maintenance and regeneration.

Here, we identify Resistin as a BMA-derived adipokine that promotes NF-κB–dependent HSC self-renewal, enhances stem cell fitness, and preserves regenerative myeloid and lymphoid hematopoiesis. We further demonstrate that RXR signaling in mature BMA is required for Resistin expression, establishing a previously unrecognized BMA– RXR–Resistin–NF-κB axis as a critical extrinsic regulator of HSC function.

## Results

### Loss of adipocyte RXRα/β does not affect steady-state hematopoiesis but impairs regenerative myelopoiesis following stress

We first evaluated the functional relevance of adipocyte RXR signaling in hematopoiesis. Murine BMA from active hematopoietic BM of young mice^15^ express Adiponectin (*Adipoq*), an adipokine associated with hematopoietic activity ^16,17^, with levels influenced by age (**Supplementary Figure 1a**). To specifically interrogate RA-independent RXR activity in adipocytes, we generated tamoxifen-inducible, adiponectin-expressing cell-specific RXRα-and RXRβ-deficient (AdipoQ-Cre-ERT2; RXRα/β^flox/flox^) mice. Cre-mediated RXR deletion was induced via intraperitoneal tamoxifen injection in both AdipoQ-Cre-ERT2; RXRα/β^flox/flox^ (Adipo-RXRα/β^Δ/Δ^) and control AdipoQ-Cre-ERT2; RXRα/β^+/+^ (WT) mice (**Figure 1a**).

**Figure 1:**
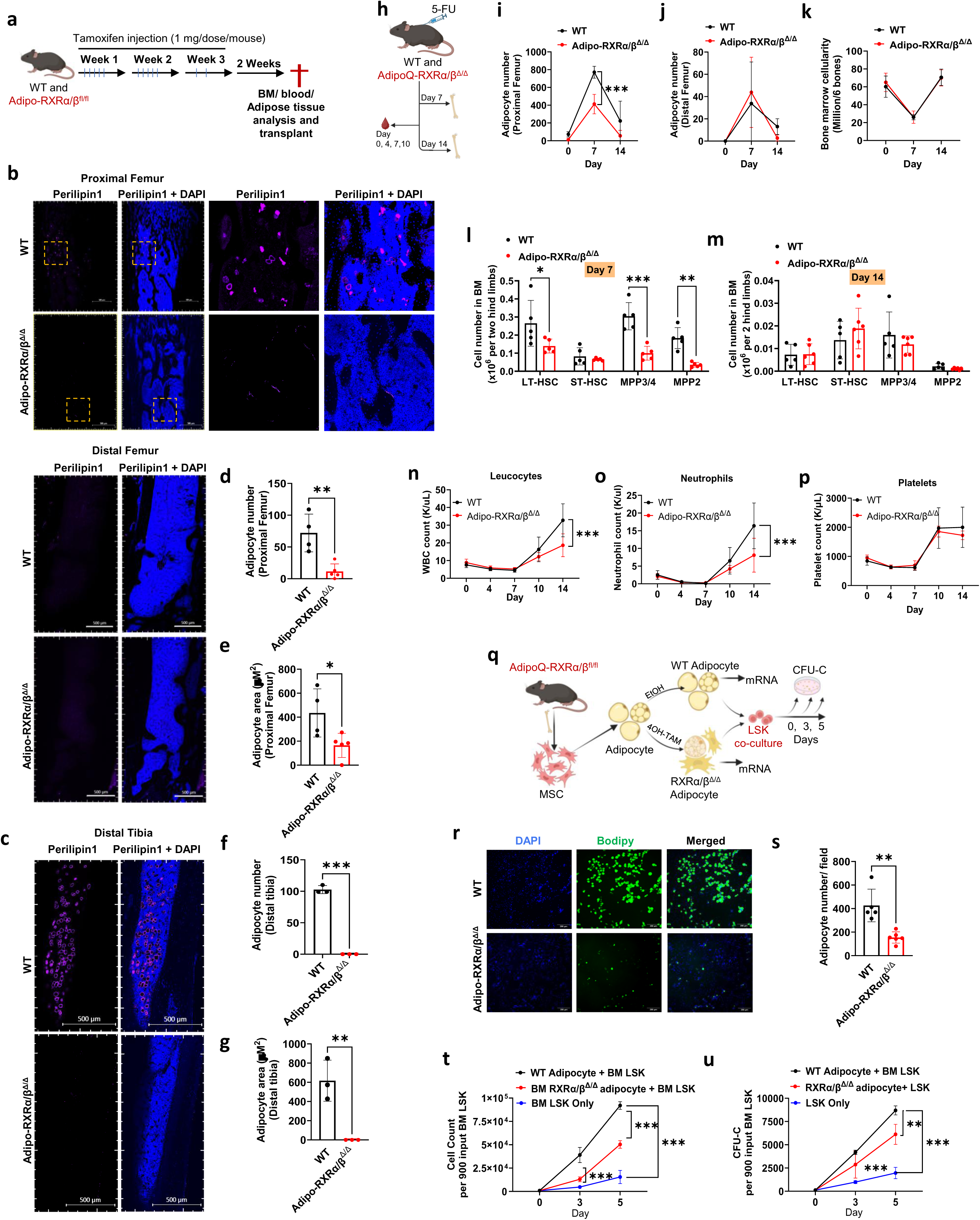
Adipocyte-specific RXRα/β deletion results in an overall decrease in adipocyte content followed by reduced hematopoietic recovery. (a) Schematic representation of tamoxifen treatment strategy before subjecting to experimental testing to induce adipocyte-specific RXRα/β deletion. (b-c) Confocal images of Perilipin1 staining of the mouse hind limb bones. Scale bars = 500 μm (d-g) Quantification of the adipocyte number and size in the bone marrow. (h) 5-FU treatment strategy. (i-j) Quantification of adipocyte numbers in the proximal and distal femur after 5-FU treatment. (k) Bone marrow cell number after 5-FU treatment. (l-m) Cell numbers of LT-HSC, ST-HSC, MPP3/4, and MPP2 in bone marrow after 5-FU treatment. (n-p) Leucocyte, Neutrophil, and platelet count in blood after 5-FU treatment. (q) Schematic representation of adipocyte differentiation and RXRαβ KO adipocyte production. (r) Fluorescence microscopic images of adipocytes grown in vitro. Scale bar=200 μm (s) Quantification of the adipocyte numbers per field. (t-u) Cell number and CFU count at indicated days after WT LSK co-culture with WT or RXRαβ KO adipocytes. Data are presented as mean ± SD. Unpaired t-test and two-way ANOVA were performed for statistical analysis. *p<0.05; **p<0.01; ***p <0.001.

At day 35 post-induction, BM, blood and and hind limb BMA were analyzed. Adipo-RXRα/β^Δ/Δ^ mice exhibited normal body weight (**Supplementary Figure 1b**) but displayed a near-complete loss of tibial adipocytes and a ∼65% reduction in both the number and size of femoral BMAs (**Figures 1b-g**). Efficiency of deletion in Adipo-RXRα/β^Δ/Δ^ mice was assessed in white adipocyte tissue (WAT) where the expression of RXRα and RXRβ were significantly decreased, with no compensatory upregulation of the isoform γ of RXR (**Supplementary Figure 1c**).

Peripheral blood (PB) analysis of Adipo-RXRα/β^Δ/Δ^ mice revealed no overt hematologic abnormalities (**Supplementary Fig. 1d**), and BM cellularity was unaffected (**Supplementary Fig. 1e**). Immunophenotypic analysis, however, demonstrated ∼30% increases in both the frequency and absolute number of short-term hematopoietic stem cells (ST-HSCs) and multipotent progenitors (MPPs) (**Supplementary Fig. 1f**), along with elevated myeloid-lineage colony-forming units (CFU-Cs; **Supplementary Fig. 1g**). Despite these changes, serial competitive repopulation assays revealed no significant impairment in the repopulating ability of Adipo-RXRα/β^Δ/Δ^ BM compared to WT controls (**Supplementary Fig. 1h**), indicating that steady-state hematopoiesis remains functionally intact in the absence of RXR in adipocytes.

We next tested whether BM stress unmasks a requirement for adipocyte RXR. Following 5-fluorouracil (5-FU) treatment, a non-lethal, transient hematopoietic stressor that induces BMA content upregulation^9^ (**Fig. 1h**), WT mice displayed a ∼70-fold expansion of proximal femoral BMA, a response that was markedly blunted in Adipo-RXRα/β^Δ/Δ^ mice (**Fig. 1i; Supplementary Fig. 2a**). In contrast, distal femoral BMA expansion was unaffected by RXR loss (**Fig. 1j**), suggesting a regional sensitivity of BM adipose tissue (BMAT) to RXR deficiency. Despite these differences, hematopoietic BM cellularity remained intact after 5-FU in Adipo-RXRα/β^Δ/Δ^ mice (**Fig. 1k**).

Functionally, Adipo-RXRα/β^Δ/Δ^ mice exhibited significantly reduced numbers of long-term HSCs (LT-HSCs), MPP3/4, and MPP2 populations at day 7 post-5-FU (**Fig. 1l**), although these deficits resolved by day 14 (**Fig. 1m**). Peripheral blood analysis revealed delayed recovery of leukocytes and neutrophils, but not platelets, in Adipo-RXRα/β^Δ/Δ^ mice (**Figs. 1n–p**). Together, these findings indicate that while adipocyte RXR is dispensable for steady-state hematopoiesis, it is required for timely regenerative myelopoiesis following cytotoxic stress.

### RXR-expressing BMA directly regulate hematopoietic progenitor expansion and HSC fitness

To determine whether BM adipocytes (BMAs) directly influence HSPC function, mesenchymal stem cells (MSCs) were isolated from WT and Adipo-RXRα/β^Δ/Δ^ BM, differentiated into adipocytes, and co-cultured with WT Lineage^−^/Sca-1^+^/c-Kit^+^ (LSK) cells (**Fig. 1q**; **Supplementary Fig. 2c**). RXRα/β-deficient BM MSCs produced significantly fewer mature adipocytes (**Figs. 1r–s**), and these cells exhibited a markedly reduced capacity to support LSK cells expansion and colony-forming unit (CFU-C) output in vitro (**Figs. 1t–u**). These findings indicate that BMAs provide direct, RXR-dependent cues that positively regulate hematopoietic progenitor activity.

To assess the in vivo relevance of RXR-expressing BMAs, we transplanted lethally irradiated WT or Adipo-RXRα/β^Δ/Δ^ congenic recipients with CD45.1^+ WT BM cells (**Fig. 2a**). Although donor chimerism in peripheral blood was comparable between groups, Adipo-RXRα/β^Δ/Δ recipients exhibited delayed leukocyte recovery (**Fig. 2b–d**). By 16 weeks post-transplant, BM cellularity and myeloid cell content were reduced by ∼50% in Adipo-RXRα/β^Δ/Δ^ recipients despite normal engraftment levels (**Fig. 2e–f**). To further test whether loss of adipocyte RXR signaling imposes lasting effects on HSPCs, we performed serial BM transplantation from primary WT or Adipo-RXRα/β^Δ/Δ^ chimeras into secondary WT hosts (**Fig. 2a**). Secondary recipients of Adipo-RXRα/β^Δ/Δ^ BM displayed persistently reduced donor chimerism in PB, BM, and spleen. Although overall BM and spleen cellularity remained intact, myeloid content in BM and lymphoid counts in spleen were reduced by ∼70% and ∼60%, respectively. These results demonstrate that RXR-expressing BMAs provide essential extrinsic cues to support hematopoietic reconstitution.

**Figure 2:**
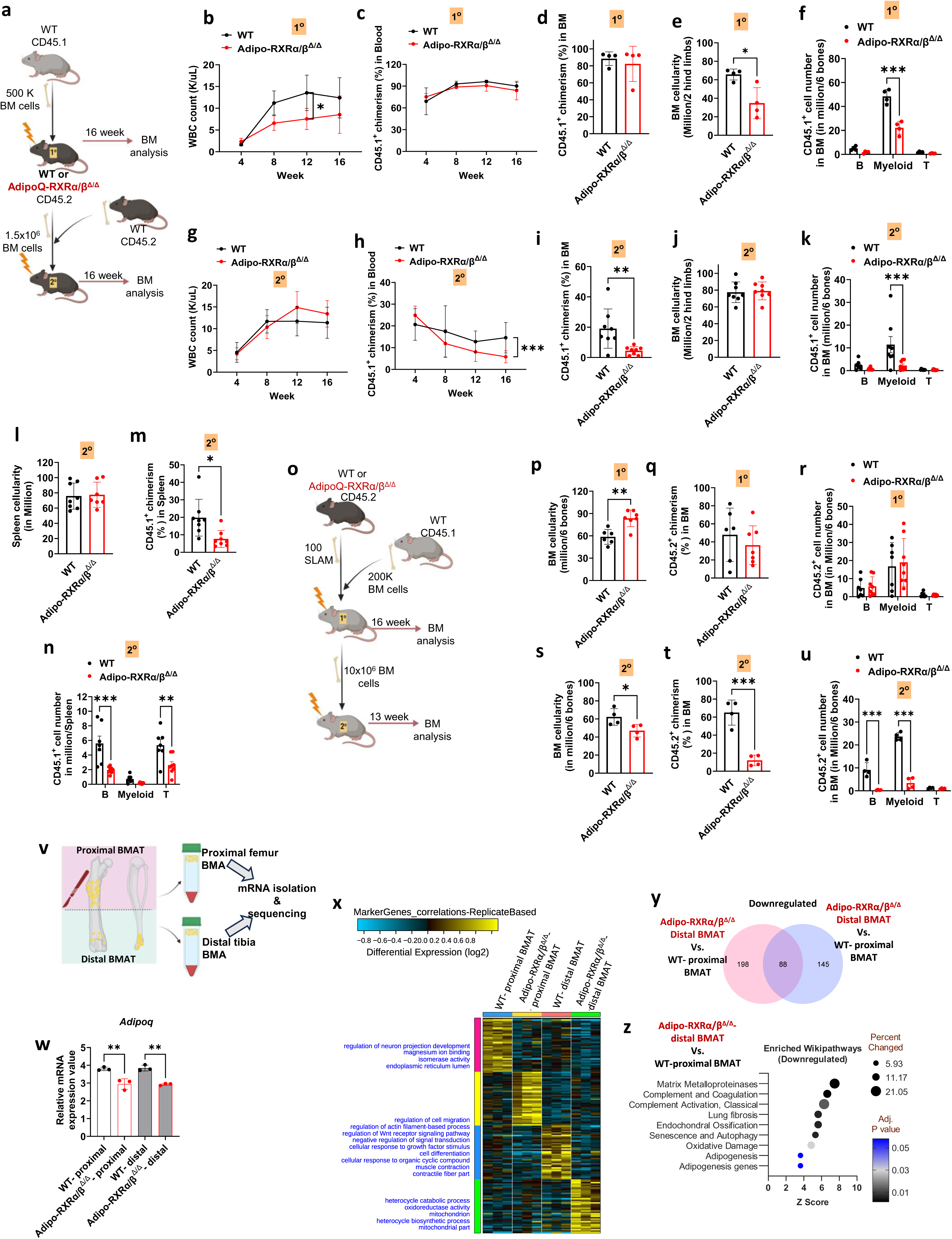
Adipocyte-specific RXRα/β expression is necessary for hematopoietic regeneration with significant changes in regulated BM adipocytes. (a) Schematic representation of WT and Adipo-RXRαβ^Δ/Δ^ mice irradiation and transplant followed by competitive secondary transplant and subsequent analysis. (b) Leucocyte count and (c) chimera analysis of primary recipients in blood. (d) Quantification of donor chimerism, (e) bone marrow cell number, (f) B, myeloid, and T cell number in primary recipients. (g) Leucocyte count and (h) Chimera analysis of secondary recipients in blood. Quantification of (i) donor chimerism, (j) Bone marrow cell number, (k) B, myeloid, and T cell number in the bone marrow of secondary recipients. Quantification of (l) total spleen cell number, (m) donor chimerism, (n) B, myeloid, and T cell number in the spleen of secondary recipients. (o) Schematic representation of serial competitive repopulation assay with isolated SLAM from bone marrow. Quantification of (p-u) Bone marrow cell number, donor chimerism, B, myeloid, and T cell number in primary and secondary recipients. (v) Schematic representation of Constitutive and Regulatory bone marrow adipocyte isolation. (w) Average (Raw) expression values of AdipoQ. (x) Heatmap of differentially regulated marker genes. (y) Venn diagram showing common differentially regulated genes between Constitutive and Regulatory adipocytes. (z) Enriched Wiki pathways of downregulated genes in Constitutive adipocytes. Data are presented as mean ± SD. Unpaired t-test and two-way ANOVA were performed for statistical analysis. *p<0.05; **p<0.01; ***p <0.001.

Because serial transplantation of whole BM from Adipo-RXRα/β^Δ/Δ mice did not reveal intrinsic HSPC defects (**Supplementary Fig. 1g**), we hypothesized that accessory BM niche cells may compensate for loss of BMA-derived signals. To directly test HSC fitness, we transplanted highly purified SLAM HSCs (LSK/CD150^+^/CD48^−^) from WT or Adipo-RXRα/β^Δ/Δ^ mice into lethally irradiated WT hosts in sequential primary and secondary competitive repopulation assays (**Fig. 2o**). Strikingly, HSCs from Adipo-RXRα/β^Δ/Δ^ mice exhibited profound self-renewal defects, with significantly reduced donor chimerism and multilineage reconstitution in both primary and secondary hosts (**Fig. 2p–u**). These findings establish RXR-dependent BMA cues as critical regulators of HSC self-renewal, fitness, and long-term hematopoietic regeneration.

To define the molecular basis of RXR-dependent BMAT function, we isolated adipocytes from the proximal vs distal femur/tibia regions (**Fig. 2v**) and performed RNA sequencing analysis. In Adipo-RXRα/β^Δ/Δ^ mice, adipocytes from both regions exhibited marked reduced expression of adiponectin *(Adipoq)* (**Fig. 2w**), suggesting a shift toward enrichment in low-adiponectin expressing BMA subsets. Transcriptomic analysis revealed 233 genes downregulated in Adipo-RXRα/β^Δ/Δ^ proximal BMAT and 286 in Adipo-RXRα/β^Δ/Δ^ distal BMAT, with only 88 genes overlapping, indicating region-specific transcriptomic regulation by RXR. Notably, no significantly enriched gene pathways with an emphasis on hematopoiesis were identified (**Fig. 2x-z; Supplemental Table 1**), suggesting that RXR acts primarily as a broad transcriptional activator of adipocyte programs in anatomically distinct BMAT regions.

### Loss of RXR signaling in BMA impairs NF-κB–associated transcriptional programs in HSCs

To define the molecular basis of defective hematopoiesis in Adipo-RXRα/β^Δ/Δ^ mice, we first performed bulk RNA sequencing (RNA-seq) on sorted HSCs. This analysis revealed a striking global transcriptional suppression, with ∼90% of genes significantly downregulated, including multiple components of TNF receptor (TNFR) and NF-κB signaling pathways critical for HSC maintenance and stress response (**Figs. 3a–c; Supplemental Table 2**).

**Figure 3:**
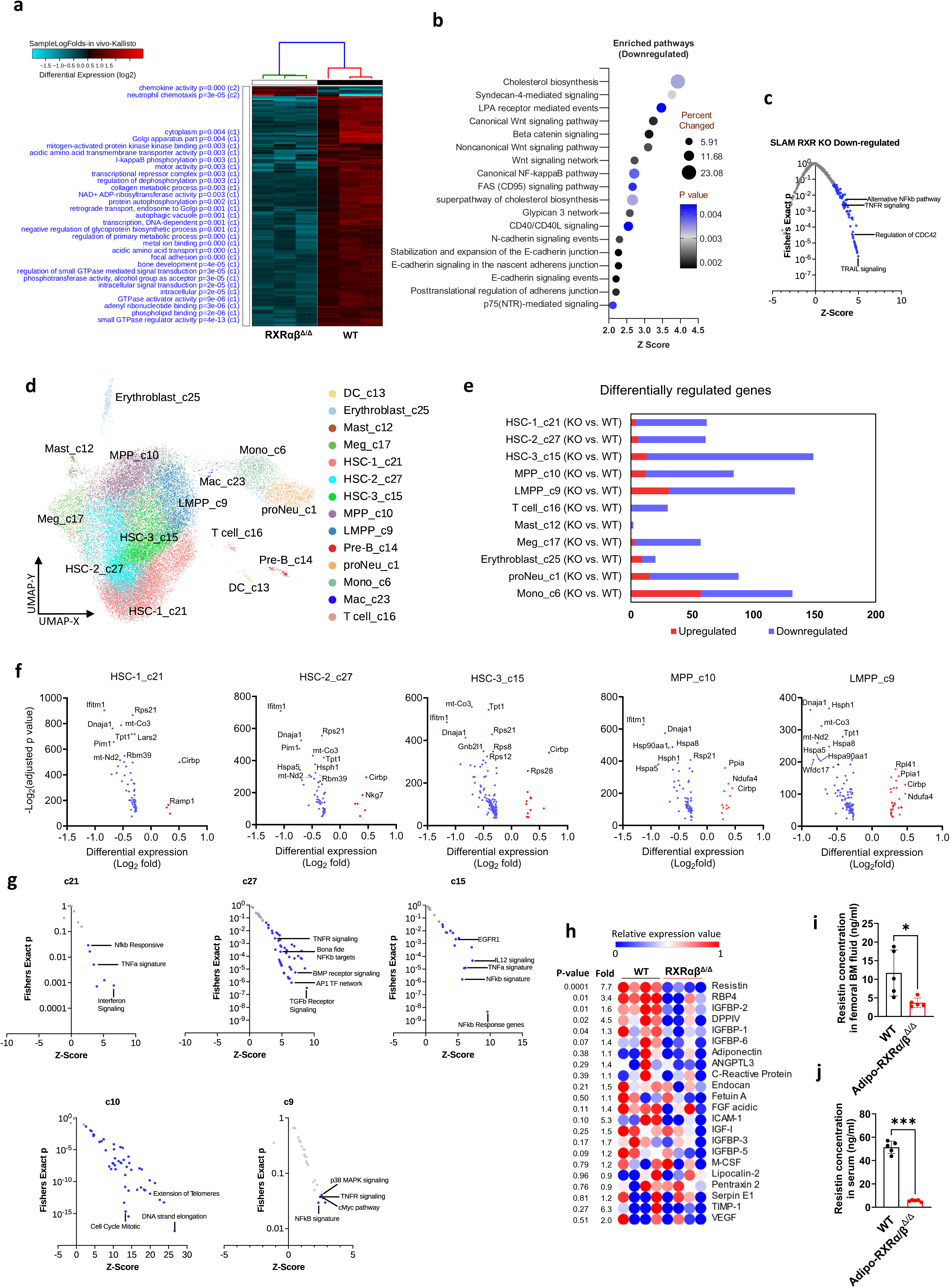
Bone marrow adipocyte-specific RXRαβ deletion reduced pro-inflammatory NF-kB signaling and Wnt pathway in the bone marrow HSCP population, and reduced Resistin in the bone marrow microenvironment. (a) Heat map showing the differentially regulated pathways in SLAMs sorted from WT and Adipo-RXRα/β^Δ/Δ^ mice. (b) Pathway analysis with downregulated DEGs (P≤0.05, FC ≤−1.5). (c) Pathway enrichment analysis of different clusters with differentially regulated genes. (d) UMAP plot with different population clusters in WT and Adipo-RXRαβ^Δ/Δ^ derived LSK cells. (e) Number of differentially regulated genes in different population clusters. (f) Volcano plot of differentially regulated genes in three different clusters of HSCP, MPP and LMPP. (g) Pathway enrichment analysis of different clusters with differentially regulated genes from scRNASeq analysis. (h) Heatmap of different adipokine levels in bone marrow extracellular fluid of WT and Adipo-RXRαβ^Δ/Δ^ mice. (i-j) Quantification of Resistin concentration in BM extracellular fluid and serum in WT and Adipo-RXRαβ^Δ/Δ^ mice. Data are presented as mean ± SD. Unpaired t-test was performed for statistical analysis. *p<0.05; **p<0.01; ***p <0.001.

To determine whether this suppression Supplemental across the progenitor hierarchy, we conducted scRNA-seq of LSK cells (**Supplemental Table 3**). Unsupervised clustering identified 14 distinct transcriptional populations (**Fig. 3d**), including three clusters corresponding to HSCs (c21, c27, c15) and two corresponding to multipotent progenitors (MPP, c10) and lymphoid-primed MPPs (LMPPs, c9)^18^. Across these five clusters, >90% of dysregulated genes were significantly downregulated in Adipo-RXRα/β^Δ/Δ^ cells (**Fig. 3e**). Only 12 genes were consistently upregulated, including ribosomal proteins (*Rpl15, Rpl27, Rpl28, Rpl35, Rpl36, Rpl37a, Rpl41*), mRNA splicing regulators (*Snrpg, Cirbp*), and mitochondrial respiratory chain components (*Atp5g2, Cox6b1, Nduf4*), suggesting selective enhancement of translational and metabolic functions within an otherwise repressed transcriptome (**Fig. 3f**).

Notably, the most strongly downregulated genes included *Ifitm1* and *Pim1*, both positive regulators of NF-κB signaling and critical mediators of HSC regenerative capacity^19,20^. Additional suppressed genes encompassed the mitochondrial respiratory chain components (*mt-Co3, mt-Nd2, mt-Atp6*), protein synthesis regulators (*Rps21, Lars2, Rps8, Rps12*), and protein-folding chaperones (*Dnaja1, Hspa1, Hspa5*), all previously implicated in NF-κB–dependent HSC maintenance and stress responsiveness^21–23^ (**Fig. 3e**). Pathway enrichment confirmed consistent suppression of TNF/NF-κB signaling across progenitor clusters, except for MPPs (c10), which showed lower pathway engagement (**Fig. 3g**).

Together, these findings demonstrate that RXR-expressing BMAs sustain HSC transcriptional integrity by maintaining tonic NF-κB signaling. Loss of BMA RXR results in broad expression of NF-κB–associated metabolic and proteostatic programs, implicating a non–cell-autonomous, BMA-dependent mechanism in preserving HSC fitness and regenerative function.

### Resistin is a BMA-derived, RXR-regulated adipokine required for maintaining NF-κB signaling in HSCs

Adipokines are adipocyte-secreted cytokines that mediate intercellular communication within the BM microenvironment (ME). To determine whether RXR signaling in adipocytes regulates adipokine secretion, we profiled BM supernatants from Adipo-RXRα/β^Δ/Δ^ and control mice. Five adipokines were significantly reduced: RBP4, IGFBP-1, IGFBP-2, DPPIV, and most notably, Resistin (**Fig. 3h**). Enzyme-linked immunosorbent assay (ELISA) confirmed a ∼90% and a ∼65% reduction of Resistin levels in BM extracellular fluid and serum, respectively, from Adipo-RXRα/β^Δ/Δ^ mice (**Figs. 3i-j**). In contrast, soluble stem cell factor (SCF), a major adipocyte-derived driver of hematopoietic regeneration^9^ , was unaffected (**Supplementary Figs. 2d**).

Resistin is a pro-inflammatory adipokine that activates NF-κB signaling through Toll-like receptor 4 (TLR4)^24^ by downregulating the inhibitory adaptor TNF-α receptor–associated factor 3 (TRAF3)^25^, thereby sustaining NF-κB pathway activation^26^. To identify the cellular source of Resistin in the BM, we employed tamoxifen-inducible Adiponectin-Cre–driven TdTomato reporter mice, which allow the distinction between mature Perilipin^+^ TdTomato^+^ mature adipocytes and Perilipin^−^ TdTomato^+^ adipocyte precursors in BM (**Supplementary Figs. 3a–b**). Resistin mRNA expression was detected exclusively in the adipocyte-rich floating fraction, with no detectable expression in the pelleted CD45^−^/TdTomato^+^ stromal cell fraction (**Supplementary Figs. 3c–e**), indicating that mouse Resistin is produced specifically by mature BM adipocytes.

Consistently, both secreted Resistin protein and *Resistin* (*Retn*) mRNA expression were significantly reduced in cultured adipocytes derived from Adipo-RXRα/β^Δ/Δ^ BM compared to WT (**Supplementary Figs. 3f–h**), confirming that RXR activity is required for Resistin expression and secretion.

Together, these results identify Resistin as a mature BMA–derived, RXR-regulated adipokine and establish it as a key extrinsic factor that sustains tonic NF-κB signaling in HSCs. Loss of Resistin in the Adipo-RXRα/β^Δ/Δ^ niche provides a mechanistic explanation for the global suppression of NF-κB–associated transcriptional programs and impaired HSC function.

### Resistin is necessary and sufficient to maintain HSC regenerative fitness following myeloablative stress

To assess the functional role of Resistin in hematopoietic regeneration, WT mice were treated with 5-FU followed by administration of either a polyclonal anti-mouse Resistin neutralizing antibody ^27,28^ with capacity to transiently abrogate Resistin signaling, or pre-immune rabbit IgG as control. BM hematopoietic cells were then harvested and subjected to serial competitive repopulation assays in lethally irradiated congenic recipients (**Supplementary Fig. 3h**). Resistin neutralization led to a significant decrease in donor-derived PB and BM chimerism, reduced BM cellularity, and impaired myelopoiesis in both primary and secondary transplants (**Supplementary Figs. 3i–q**). Immunophenotypic analysis of BM cellularity revealed a marked reduction in the content of LSK cells including long-term (LT)-HSCs, short-term (ST)-HSCs, and MPP3 cells in secondary recipients (**Supplementary Figs. 3r–u**), whereas MPP2 and lymphoid-biased MPPs (LMPPs) were unaffected (**Supplementary Figs. 3v–w**). These data indicate that endogenous Resistin is essential for HSC expansion and MPP3-driven myeloid regeneration following hematopoietic injury.

Resistin is a cysteine-rich hormone, expressed by murine adipocytes and human macrophages^29^, that circulates in both low molecular weight (LMW; monomer/dimer) and high molecular weight (HMW; trimer/hexamer) forms^30^. To test which isoform supports HSC regenerative capacity, purified WT BM LSK cells were treated ex vivo for 16 hours with vehicle, E. coli–derived recombinant LMW Resistin (monomers/dimers), or NS0 myeloma–derived HMW hexameric Resistin at concentrations of 10 or 100 ng/mL prior to transplantation into lethally irradiated mice, followed by transplantation into lethally irradiated recipients and serial repopulation analysis (**Fig. 4a**). Vehicle-treated LSK cells lost competitive repopulating capacity by tertiary transplantation. In contrast, LSK cells pretreated with LMW Resistin displayed resistance to exhaustion, maintaining high donor-derived PB chimerism and multilineage (myeloid, B, and T cell) output in tertiary recipients (**Fig. 4b–e**). BM analysis confirmed enhanced donor-derived reconstitution and myelopoiesis (**Figs. 4f–k**) with no significant effect on total BM cellularity **(Supplementary Fig. 4a)**. By comparison, HMW Resistin, produced only modest gains in secondary recipients at molar equivalent concentrations (100 ng/mL; **Supplementary Figs. 4b–h**), suggesting that LMW Resistin possess superior biological potency in supporting durable HSC function.

**Figure 4:**
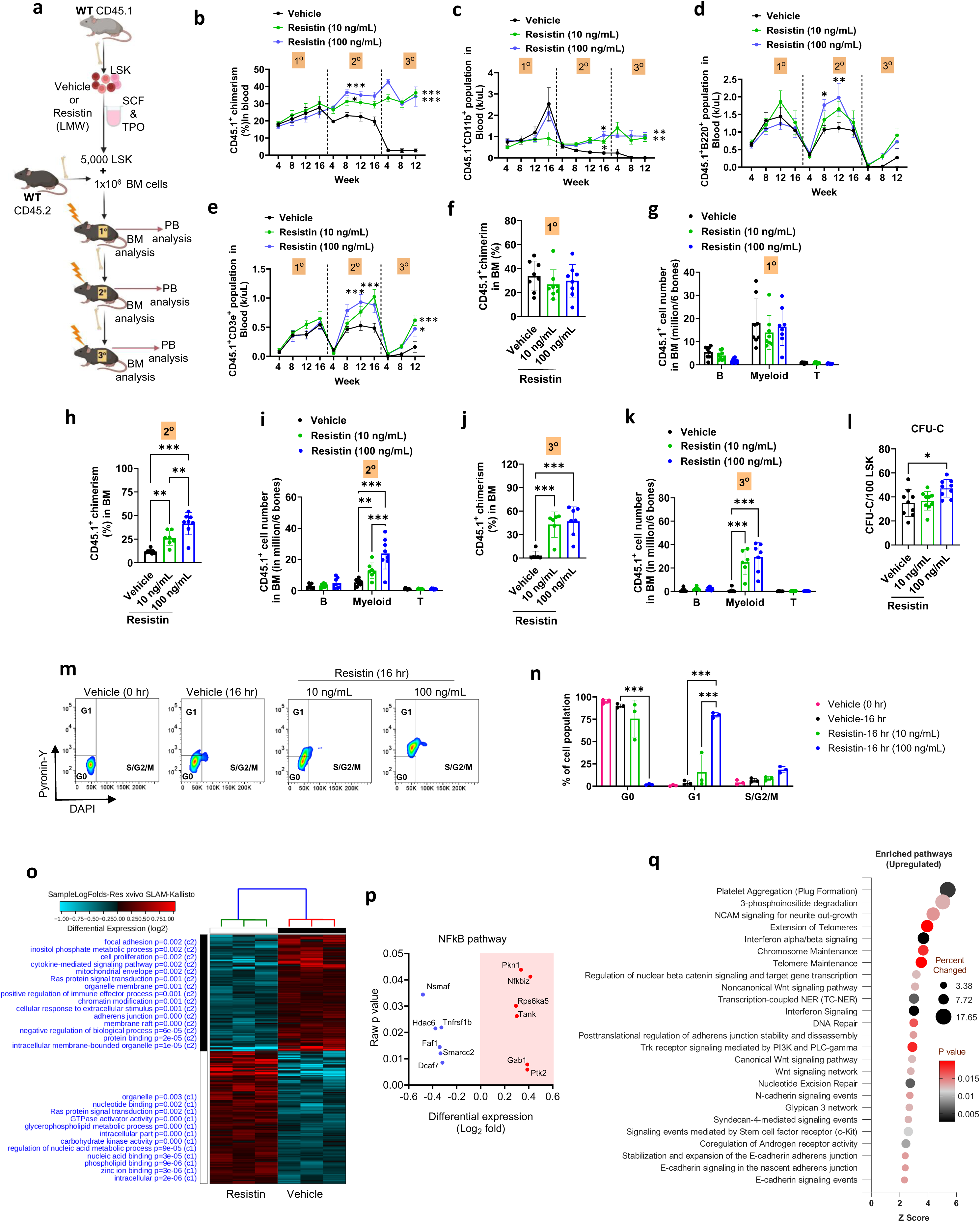
Resistin is sufficient for long term hematopoiesis without compromising lymphocyte production. (a) Schematic representation of ex vivo Resistin treatment to WT LSK cells and serial competitive repopulation assay. (b-e) Quantification of blood chimera, myeloid, B and T cell repopulation after serial competitive transplant. (f-k) Quantification of bone marrow chimera; B, myeloid and T cell number after serial competitive transplant. (l) Quantification of CFU-C after ex vivo Resistin treatment to WT LSK. (m) Flowcytometric plot for Pyronin Y-DAPI staining. (n) Quantification of cell frequency in different cell cycle stages. (o) Heat map showing the differentially regulated pathways in SLAMs sorted from WT and Adipo-RXRα/β^Δ/Δ^ mice. (p) DEGs related to NF-kB signaling. (q) Pathway analysis with upregulated DEGs (P≤0.05, FC ≤−1.2). Data are presented as mean ± SD. Unpaired t-test, two-way ANOVA and chi square test were performed for statistical analysis. *p<0.05; **p<0.01; ***p <0.001.

To further define the action of LMW Resistin on progenitors, we assessed its effect on myeloid CFU-C activity. LMW Resistin induced only a marginal increase in CFU-C formation (**Fig. 4i**), supporting that HSCs rather than myeloid progenitors are the primary target of LMW Resistin. Cell cycle analysis of HSCs treated with LMW Resistin revealed an accelerated transition into G1 phase (**Figs. 4m–n**), consisting with a role in promoting proliferation while preserving stemness. To elucidate the molecular pathways involved, RNA sequencing was performed on sorted HSCs treated with 10 ng/mL LMW Resistin. Transcriptomic profiling revealed differential regulation of NF-κB signaling (**Figs. 4o–p; Supplemental Table 4**) and enrichment of pathways associated with Wnt signaling, telomere maintenance, interferon response, c-Kit signaling, DNA repair, and junctional signaling (**Fig. 4q**). Together, these results demonstrate that LMW Resistin is both necessary and sufficient to promote HSC proliferation, preserve long-term regenerative capacity, and activate key signaling pathways that support HSC fitness and myeloid regeneration under stress conditions.

### Resistin promotes NF-κB–dependent HSC self-renewal and long-term hematopoiesis

NF-κB signaling plays a critical role in regulating HSC homeostasis and hematopoiesis^22^. Consistent with this, inflammatory NF-κB pathway components were transcriptionally downregulated in HSCs derived from Adipo-RXRα/β^Δ/Δ^ mice. To test whether Resistin regulates NF-κB activity, we treated WT HSCs with monomeric LMW Resistin, which significantly enhanced NF-κB nuclear translocation in WT HSCs (**Supplementary Figs. 4j, k**).

To directly assess the functional consequences of Resistin–NF-κB signaling on HSC fate, we performed c-Myc polarity and paired-daughter division assays. Resistin treatment significantly increased the frequency of symmetric self-renewing divisions, indicative of enhanced self-renewal capacity, wherea the pre-treatment with two mechanistically distinct NF-κB pathway inhibitors (QNZ and BAY 11-7082) abrogated this effect (**Figs. 5a-e**). These findings demonstrate that Resistin promotes HSC self-renewal through NF-κB–dependent mechanisms.

**Figure 5:**
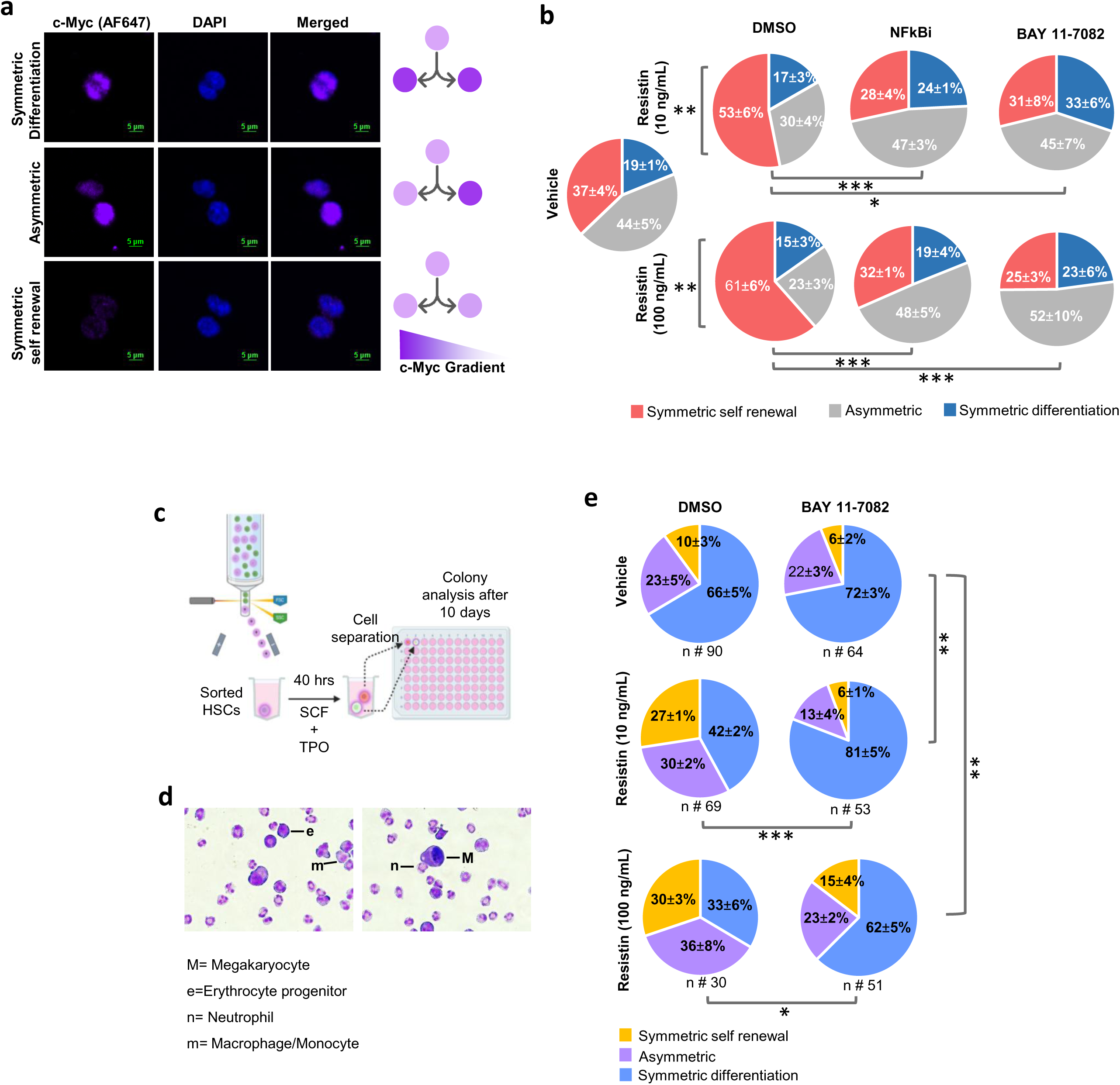
Resistin supports hematopoietic self-renewal through the NF-kB signaling pathway. (a) Confocal microscopic images of the first HSC division with c-MYC staining along with (b) quantification of the frequency of asymmetric, symmetric differentiation, and symmetric self-renewal divisions. (c) Schematic representation of paired daughter cell assay. (d) Microscopic images of colonies stained with QUICK-diff. (e) Quantification of the frequency of asymmetric, symmetric differentiation, and symmetric self-renewal divisions. Scale bar = 5 μm. Data are presented as mean ± SD. Unpaired t-test, two-way ANOVA and chi square test were performed for statistical analysis. *p<0.05; **p<0.01; ***p <0.001.

Together with our transplantation data, these results identify BMA-derived Resistin as a positive regulator of HSC function that promotes HSC long-term function. By activating NF-κB, Resistin, preserves HSC fitness across serial transplantats, and maintains long-term balanced myeloid and lymphoid hematopoiesis following stress-induced regeneration.

## Discussion

BM HSCs depend on environmental signals from the BM ME for their maintenance and regenerative capacity, particularly following myeloablation. BMAs, which originate from MSCs and share a common origin with other stromal cell types, possess distinct phenotypic and functional properties compared with other well characterized tissue adipocytes. In addition to their established roles in bone health and metabolism, BMAs are closely associated with HSCs and hematopoietic progenitors^31,32^.

Traditionally, BMAs have been considered negative regulators of hematopoiesis, based on their expansion in aged BM^33–35^ and the association of adipocyte-rich regions with myeloid biased hematopoietic activity^35^, this paradigm has been challenged by studies demonstrating context-dependent regulatory roles, both positive and negative^9^.

Notably, BMAs are rare in the marrow of young adult mouse bones but expand markedly following myeloablation^9^, particularly in the long bones of the hind limbs, which harbor both constitutive and regulatory adipocyte populations ^36–38^. Irradiation not only depletes HSCs but also damages their niche by destroying sinusoidal blood vessels and depleting stromal cells^39–41^. Niche regeneration is essential for hematopoietic recovery^42,43^ and niche BMAs seem to contribute to supporting hematopoietic recovery^9,44^. Signaling by retinoic acid (RA), locally synthesized by the regenerating BM niche cells, plays a central role in the BM hematopoietic response to stress^45–48^. RA signals through nuclear receptors, including Retinoid X Receptors (RXRs), which act as homo- or heterodimers or heterodimers to activate transcriptional programs in HSC^14,49^, and are also required for adipocyte differentiation via PPARγ^50^. Based on these insights, we hypothesized that RXR signaling in BMAs positively regulates hematopoiesis by controlling the expression and secretion of adipokines with functional relevance to HSC maintenance and regeneration.

Our study identifies RXR-mediated expression of the adipokine Resistin in BMAs as essential for activating NF-κB signaling in HSCs. Through gain- and loss-of-function approaches, in vivo and ex vivo, we demonstrate that BMA-derived Resistin functions as a selective and potent regulator of HSC regeneration, capable of overriding suppressive environmental cues via NF-κB–dependent signaling. This cascade enhances HSC self-renewal, functional fitness, and multilineage hematopoietic regeneration.

Using tamoxifen-inducible Adipo-RXRα/β^Δ/Δ^ mice, we found that RXR deficiency in adipocytes does not perturb steady-state hematopoiesis but profoundly impairs hematopoietic regeneration following 5-fluorouracil (5-FU)–induced injury. Functionally, RXR-deficient BMAs display reduced expression of adiponectin and Resistin diminished capacity to support HSC expansion in vitro, and compromised HSC self-renewal and fitness in vivo.

BMAs are an important source of SCF, a driver of hematopoietic regeneration ^9^. Although we cannot rule out the potential role of deficient transmembrane SCF ^51^ in Adipo-RXRα/β^Δ/Δ^, interestingly, we did not observe a significant alteration in the levels of soluble SCF in WT and BMA RXR deleted mouse bone marrow extracellular fluid, suggesting that adipocytic SCF expression is RXR independent.

Mechanistically, RXR signaling operates as a transcriptional rheostat in BMAs, modulating not only their adipogenic program but also their secretome, including key hematopoietic niche factors. Transcriptomic analyses of RXR-deficient HSCs reveal broad downregulation of NF-κB– and TNFR-related gene signatures, consistent with a loss of Resistin-driven signaling input. Resistin is secreted exclusively by mature, predominantly regulated BMAs, which are enriched in the proximal femurs and tibiae, and its expression is drastically reduced in Adipo-RXRα/β^Δ/Δ^ mice. Functional assays demonstrate that low molecular weight (LMW) Resistin restores HSC fitness and regenerative capacity, promotes NF-κB activation, and enhances symmetric self-renewing divisions in an NF-κB–dependent manner, in both in vitro and in vivo settings. Conversely, neutralization of Resistin or inhibition of NF-κB signaling abrogated these regenerative effects, confirming the critical role of this axis in hematopoietic recovery.

Interestingly, while tonic NF-kB activation is HSC protective as evidenced in in vivo experiments of NF-kB loss-of-function^52^, the expression of a constitutively active form of IKK2 in HSCs in mice results in a the opposite phenotype, compromising the pool size and activity of BM HSC ^53^, suggesting that the effect of Resistin on NF-kB signaling is of tonic nature.

Overall, this study defines a novel RXR–Resistin–NF-κB signaling axis in BMAs as a critical extrinsic regulator of HSC function. This pathway ensures the tonic inflammatory signaling necessary for long-term HSC self-renewal and balanced multilineage hematopoietic regeneration. These insights underscore adipocyte-derived Resistin as a central mediator of BM regenerative capacity and point to BMA-intrinsic signaling as a therapeutic target for enhancing hematopoietic recovery after myeloablation.

## STAR methods

### Mice and inducible RXRα and RXRβ deletion

To induce adipocyte-specific deletion of *Rxrα* and *Rxrβ*, 8–10-week-old AdipoQ-CreERT2; Rxrα ^fl/fl^; Rxrβ^fl/fl^ and littermate Rxrα^fl/fl^; Rxrβ^fl/fl^ (WT) controls were administered tamoxifen (Sigma, St. Louis, MO; Cat #T5648) dissolved in corn oil (20 mg/mL), by injecting 1 mg Tamoxifen/animal (10 mg/ml) per dose during consecutive 5 days (first week), every 3 days for other two weeks. Mice deleted for adiocytic RXRα and RXRβ (AdipoQ-RXRαβ^Δ/Δ^) were euthanized at 2 weeks after the last dose of tamoxifen. Male and female mice were studied from 2 to 6 months of age. For experiments under homeostatic conditions, untreated age- and sex-matched littermate controls were used. All animal protocols were approved by the Institutional Animal Committees of Cincinnati Children’s Hospital Medical Center and Dana-Farber Cancer Institute. B6.SJL^PtprcaPepcb/BoyJ^ (CD45.1^+^) and C57BL/6J mice were purchased from either in-house breeding facility (CCHMC) or The Jackson Laboratory (Strains #:002014 and 000664).

### 5-FU administration

Mice were injected 200 mg/kg of 5-FU (5-Fluorouracil, Fresenius Kabi, # 101710) intraperitoneally. PB samples were obtained at indicated days by retro-orbital bleeding using heparin coated capillary tubes in a EDTA coated collection tube. Absolute blood counts were performed using Hemavet (Drew Scientific) or HT-5 (Heska) CBC analyzer. Hind limb bones were harvested for either microscopy or flowcytometry analysis at indicated time points.

### Bone marrow harvest and serial competitive repopulation assay

Femurs, tibiae and two hemi-pelvic bones (6 bones) were excised from mice and crunched in PBS using a tissue mortar and filtered with 100 µM nylon filter. RBC lysis was performed with brief exposure to Pharm Lyse Buffer (BD, San Jose, CA; Cat # 555899). BM cellularity is presented as the count of BM cells after RBC lysis of these six bones. For total bone marrow (BM) transplantation, equal amounts of CD45.2^+^ BM cells (WT and AdipoQ-RXRα/β^Δ/Δ^, control IgG and Anti-Resistin antibody treated mice) were mixed with congenic CD45.1^+^ B6.SJL^PtprcaPepcb/BoyJ^ competitor BM cells at 1:1 ratio and then transplanted into lethally irradiated (7+4.75 Gy, split dose) B6.SJL^PtprcaPepcb/BoyJ^ recipients. The cell mixing ratio was verified by analyzing the CD45.2 and CD45.1 percentage through flow cytometry. For secondary transplantation experiments, BM cells were pooled from each recipient for each group and serially transplanted into secondary recipients (10×10^6^ cells/mouse). This process was terminated after 16-20 weeks of secondary or tertiary reconstitution, and the mice were euthanized for analysis.

In some experiments, HSC (SLAM, Lin^-^/c-kit^+^/Sca1^+^/CD150^+^/CD48^-^) were sorted from WT and AdipoQ-RXRab^Δ/Δ^ BM and 100 SLAM cells were transplanted along with 200,000 congenic CD45.1^+^ B6.SJL^PtprcaPepcb/BoyJ^ competitor BM cells into lethally irradiated (7+4.75 Gy, split dose) B6.SJL^PtprcaPepcb/BoyJ^ recipients.

Peripheral blood was collected via retro-orbital bleeding, and blood chimera of primary, secondary, and tertiary recipients was measured at indicated time points. Absolute blood counts were recorded using Hemavet (Drew Scientific, Dallas, TX) or HT-5 (Heska, Loveland, CO) complete blood count analyzers. After RBC-lysis, the cell suspension was incubated with Fc-receptor blocker (Thermo-Fisher, #14-9161-73, Waltham, MA) antibody. Then staining was performed by using monoclonal anti-mouse lineage antibodies against mouse CD45.2, CD45.1, CD45R (B220), CD11b and CD3ε. At the endpoint of the primary transplant, bone marrow transplanted mice were euthanized, and PB, BM, and spleen were analyzed for total CD45.2 chimera, lineage reconstitution/ absolute lineage positive cellularity, and HSC/P.

### Adipokine profiling

Adipokine profiling was performed according to the manufacturer’s protocol (R&D Adipokine kit, # ARY013) using 50 μL of bone marrow extracellular fluid. After developing the membrane, the intensity of each spot was measured by ImageJ. Heatmap was created using Morpheus software (https://software.broadinstitute.org/morpheus) with the relative intensity measurements.

### BM Adipocyte Isolation

Femurs and tibias were first crunched in PBS and the cell solution was transferred to a 50 ml tube without filtration. After a brief centrifugation, the cell solution was kept undisturbed in room temperature for 10 mins to allow enough time for the adipocytes to float. Then around 300 µL of surface layer containing floating adipocytes were carefully aspirated with a 1mL pipet tip. This adipocyte solution was then further processed for RNA isolation. For isolation of proximal and distal long bone BMAT separately, the femur and tibia were cut horizontally with a scalpel (**Figure 2v**) and crunched for BMA isolation.

### Bone marrow adipocyte analysis with microscopy

For bone marrow adipocyte imaging, freshly isolated bones (Femur and tibia) were fixed with 4% PFA for 2 days at 4 °C in a gentle shaking condition. After that, the bones were fixed in Tissue-Tek OCT Compound (Sakura Finetek, #4583) and shaved half longitudinally to expose the bone marrow with cryostat. The bones were again fixed briefly with 4 % PFA and stained with anti-Perilipin1 antibody (Invitrogen, #PA5-18694) overnight. After that, the bones were washed and stained with anti-goat AF647 tagged Secondary antibody (Invitrogen, # A-31573) for 1 hr. The bones were counterstained with DAPI. Confocal images of the bones were captured with Nikon confocal microscopy (20X objective) and stitched together. IMARIS 10.2 software was used for adipocyte detection, number, and size analysis. For adipocyte detection, surface application was used for marking the individual adipocytes. Further, the selections were manually tuned for accurate assortment of adipocytes inside the bone marrow.

For imaging the differentiated adipocytes from isolated BM-MSCs in vitro, cells were stained with Bodipy (Invitrogen, #D3922) and DAPI (Invitrogen, #62248) and imaged with Nikon wide field microscope (10X objective). Images were processed with NIS Elements software. For adipocyte counting per field, FIJI (ImageJ) software was used.

### BM-MSC isolation and adipogenic differentiation

Mouse BM-MSC was isolated following previously published protocol ^54^. Briefly, femurs, tibiae and pelvis were isolated from AdipoQ-RXRαβ^Δ/Δ^ mouse and bones were crunched and bone fragments were digested with PBS+ 0.1% BSA+ Collagenase-II 2mg/ml and Dispase 1mg/mL. After that cell suspension was washed with PBS and MSCs were grown using MesenCult Expansion Kit (Mouse) (Stemcell technologies, # 05513). Adipocyte differentiation was performed using MesenCult Adipogenic Differentiation Kit (Stemcell technologies, #05507). To induce RXR deletion, MSCs or MSC derived adipocytes were treated with 1µM of 4-OH Tamoxifen (Stemcell technologies, #74052) every 24 hours for 3 consecutive days with replenishing half of the media. On day 3, media was collected to analyze secreted Resistin levels, and the cells were used for mRNA isolation.

### Adipocyte-LSK co-culture

WT and AdipoQ-RXR KO adipocytes were generated from isolated BM-MSCs in vitro as discussed above. After that, LSKs were sorted from WT mice, and 900 cells/well were co-cultured in a 96-well plate. Cells were harvested at the indicated time points, and 300 cells/dish were plated for the CFU-C assay.

### Bone marrow extracellular fluid and serum isolation

One femur per animal was flushed out using 180 μL PBS with a 21 G needle. Then the supernatant was collected after centrifugation. The extracellular fluid was used for adipokine profiling and ELISA assays. Peripheral blood from each mouse was collected through heart puncture or retro-orbital bleeding. For retro-orbital bleeding non-heparin coated capillary tubes (Fisherbrand, #22-362-566) were used to collect the blood in 1.5 mL Eppendorf tubes to allow clotting. After clotting, the blood was centrifuged, and the serum was collected. Serum was used for ELISA assays.

### Resistin enzyme-linked immunosorbent assay (ELISA)

Resistin ELISA was performed following the manufacturer’s protocol (ABCAM; #ab205574) using blood serum or plasma and bone marrow extra-cellular fluid.

### esistin Neutralization

For Resistin neutralization, WT C57BL/6J mice were treated with intravenous anti-Resistin antibody (100 µg/mouse) (Millipore-Sigma, #AB3708) as published previously^27,55^. This rabbit polyclonal antibody was generated against a synthetic peptide from mouse Resistin (aa 83-91). Control group mice were treated with purified Rabbit IgG antibody (100 µg/mouse) (Invitrogen, Waltham, MA; Cat #02-6102) followed by the same day 5-FU (Fresenius Kabi, Lake Zurich, IL, Cat # 101710) injection intraperitoneally. After 14 days, the bone marrow from 2 hind limbs was harvested, pooled, and serial competitive transplant was performed.

### Recombinant Resistin treatment

Recombinant Resistin was purchased form R&D Systems (Minneapolis, MN). The monomer and disulfide-linked homodimer (low molecular weight) form of Resistin was produced in *E. coli* (Ser21-Ser114) (RnD # 1069-RN/CF). The homohexamer (high molecular weight) (RnD # 5335-RN-050) form of protein was sourced from mouse myeloma cell line, NS0 using the same cDNA sequence.

For ex-vivo Resistin treatment, FACS sorted WT (B6.SJL^PtprcaPepcb/BoyJ^ or C57BL/6J) LSKs from 2 hind limbs were incubated overnight (16 hrs) with or without 10 or 100 ng/ml of Resistin in presence of rmSCF (100 ng/ml, Prospec, Rehovot, IL; Cat # CYT275) and rhTPO (100 ng/ml, Prospec, Cat # CYT-346) in X VIVO-15 (LONZA, Portsmouth, NH; Cat # 02-053Q) media in U bottom 96 well plate (10K LSK/well). Cells were cultured in 37 °C, 5% CO_2_ in 100% humidified incubator. 5K LSK/mouse was transplanted with 1×10^6^ competitor congenic bone marrow cells in lethally irradiated congenic mouse.

### Colony-forming-cell assays

Mouse hematopoietic stem and progenitor cells were grown in methylcellulose medium supplemented with cytokine mixtures (Methocult GF M3434; Stem Cell Technologies, # 03434) in a 37° C incubator with 5% CO_2_ and 100% humidity. For bone marrow cells, 4k cells/dish was plated, for sorted LSK cells treated with Resistin, 100 cells/dish was plated and for sorted LSK cells co-cultured with adipocytes, 300 cell/well were plated. Colony-forming progenitors were scored on days 7-10.

### Flow cytometry and Immunophenotypic analysis of HSC/P

Bone Marrow cells were stained using a mixture of biotin-conjugated monoclonal anti-mouse lineage antibodies against CD45R (B220), Gr1 (Ly6G, Clone RB6-8C5), Mac-1 (CD11b, CloneM1/70), CD3ε (Clone 145–2C11), and TER119 (Ly-76) (all from BD). In a subsequent labeling step, the cells were incubated with a combination of streptavidin-allophycocyanin (APC)-Cy7 (BD), anti-mouse Sca-1 (Ly6A/E), anti-mouse CD117 (c-kit), anti-mouse CD16/32, anti-mouse CD48, anti-mouse CD34, anti-mouse CD135, and anti-mouse CD150 antibodies. Analysis of bone marrow nuclear cells harvested from all genotypes were immunophenotypically defined by differential expression of cell surface antigens: Lin^-^ CD45.1^+/-^ CD45.2^+/-^ C-kit^+^ Sca-1^+/-^ CD135^+/-^ CD34^+/-^ CD48^+/-^ CD150^+/-^ CD16/32^+/-^.

### Cell sorting

For sorting of LSK or SLAM population, bone marrow cells were depleted for lineage cells using Lineage depletion kit (Miltenyl Biotec, # 130-110-470). Afterwards, cells were stained with Lineage markers, SCA-1 and c-Kit for LSK sorting. Cells were stained with Lineage markers (BD, # 559971), SCA-1 and C-Kit, CD48 and CD150 anti-mouse antibodies for SLAM (LSK-CD48^-^CD150^+^) sorting. For sorting TdTomato^+^ (AdipoQ^+^) bone marrow stromal cells, the bone marrow cells were isolated and depleted for CD45^+^ cells using CD45 MicroBeads (Miltenyl Biotec, 130-052-301). Sorting was performed with FACSAria II (BD, Biosciences), and SONY MA900.

### Cell cycle analysis with DAPI-Pyronin Y

SLAMs were sorted from WT mouse and incubated with or without Resistin for 16 hrs. After that the cells were fixed with BD Fix-Perm solution (BD, #51-2090KZ) for 10 min, washed with Perm/Wash buffer (BD, #51-2091KZ), and stained with DAPI (500 nM)/ Pyronin Y (2.5 µg/mL) solution for 30 min at 37 °C. After washing, the cells were analyzed through flow cytometry.

### NF-kB cellular localization

Twelve-well chamber µ-slides (iBiDi, # 81201) were coated with RetroNectin (Takara Bio, Waltham, MA, #CH-296) for 1 hr in 37 °C incubator. Sorted SLAMs (from WT C57BL/6J mouse) were cultured in these slides with X-VIVO 15 (LONZA, # 02-053Q) containing mouse stem cell factor (SCF, 100 ng/mL, Prospec, #CYT275), Thrombopoietin (TPO, 100 ng/mL, Prospec, #CYT-346) and either with vehicle (0.1% protease free BSA (Sigma, # A7030) in PBS) or different concentration (10 ng/mL and 100 ng/mL) of Resistin for 15 mins at 37 °C, 5% CO_2,_ 100% humidified incubator. After that the cells were fixed with 4% PFA (paraformaldehyde, Electron microscopy sciences, Hatfield, MA; #15710) and permeabilized with 0.1% TRITON X-100. Then the cells were incubated with 5% goat serum and then incubated with anti-NF-kB antibody (Santa Cruz, clone C-20, #sc-372) overnight at 4 °C. Then, the cells were washed and incubated with anti-rabbit secondary antibody (Invitrogen, AF488 # A-11008 or AF647, #A-31573) in 5% goat serum for 1 hr at room temperature. Finally, cells were counter stained with 10 μg/mL DAPI (ThermoFisher, Cat # D1306) for 5 min in room temperature. Samples were mounted using Prolong Diamond antifade mounting medium (Invitrogen, # P36961). Z-stacks were acquired with Nikon A1R inverted confocal microscope or Zeiss 980 confocal microscope. The NF-kB nuclear localization was analyzed with IMARIS. The nucleus was marked with surface application and the NF-kB signals were converted into spots. Then the localization statistics of each spot were analyzed.

### Paired daughter cell assay

For sorting of SLAM population, bone marrow cells were depleted for lineage cells using Lineage depletion kit (Miltenyl Biotec, # 130-110-470). Cells were stained with Lineage markers, SCA-1 and C-Kit, CD48 and CD150 anti-mouse antibodies for SLAM (LSK-CD48^-^CD150^+^) sorting. After that the cells were either treated with DMSO or BAY 11-7082 (300 nM, Santacruz Biotechnology, #sc-200615B) for 30 mins and washed twice before sorting. First the SLAMs were sorted in bulk. From that bulk sorted SLAMs, single SLAMs/well were sorted in 96 well plates containing 20 μL of X-VIVO 15 medium supplemented with SCF (100 ng/mL), TPO (100 ng/mL) with or without Resistin (10 ng/mL and 100 ng/mL). The plates were kept in 37° C for 40 hrs to allow 1 division of the SLAMs. After that the SLAMs were separated and put in 2 different well containing IMDM medium supplemented with 10% batch selected FBS (Hyclon-Cytiva), SCF, TPO, IL-3 (20 ng/mL each), EPO (4U) and G-CSF (10 ng/mL). After 10 days, colony types were analyzed the frequency of symmetric self-renewal, symmetric differentiation and asymmetric division frequency was calculated. The colonies were mounted in a slide using cytospin and differentially stained with HemaDiff Rapid Stain Set (StatLab, #11970-16). The images were acquired with Nikon ECLIPSE Ts2.

### HSC first division fate analysis

HSC first division fate analysis with Myc staining was performed as published earlier ^56^. Sorted SLAMs (from WT C57BL/6 mouse) were cultured onto RetroNectin (TAKARA, #CH-296) coated 12-well chamber µ-slide (iBiDi, # 81201) with X-VIVO 15 (LONZA, #02-053Q) containing mouse stem cell factor (100 ng/mL, Prospec, #CYT275), Thrombopoietin (100 ng/mL, Prospec, #CYT-346) and either with vehicle control or different concentrations of Resistin for 16 hrs at 37 °C, 5% CO_2_. For NF-kB inhibitor treatment, the cells were pretreated with BAY 11-7082 (300 nM, Santacruz Biotechnology, #sc-200615B) or NF-kB activation inhibitor (10 nM, Sigma-Aldrich, #481406) for 1 hr in 37 °C, 5% CO_2_, then washed and treated with or without Resistin for 16 hrs. Nocodazole (10 nM, Sigma, #M1404) was added, and cells were cultured for an additional 24 hours and then fixed with 4% paraformaldehyde (PFA) (Electron Microscopy Sciences, #50-980-487). Samples were permeabilized with 0.1% TRITON X-100 (Sigma-Aldrich, #T8787) for 10 minutes and then blocked with 5% goat serum (Sigma, # 50062Z) in PBS for 1 hour in RT. Samples were labelled with anti-MYC Rabbit polyclonal Ab (Cell signaling, Danvers, MA , # 5605) overnight at 4 °C. Then, the cells were incubated with anti-rabbit secondary antibody (Invitrogen, AF488 # A-11008 or AF647, #A-31573) in 5% goat serum for 1 hr at 4 °C. Finally, cells were counterstained with 10 μg/mL DAPI (ThermoFisher, # D1306) for 5 min in room temperature. Samples were mounted using Prolong Gold antifade mounting medium (Invitrogen, #P36930). Images were taken with an LSM 800 confocal microscope system (Zeiss) using a 20X/25X objective lens and analyzed by ZEN software.

### Quantitative real time PCR (Q-PCR)

Total mRNA was isolated from following the manufacturer’s protocol (Qiagen, #74104) following manufacturer’s instructions, and cDNA was prepared using Taq Man reverse transcription reagent (Applied Biosystems, Life technologies, catalog N8080234). The mRNA expression levels of GAPDH, RXRα, RXRβ, and Resistin were analyzed by Q-RT-PCR (qRT-PCR) assay using TaqMan Universal PCR master mix and gene-specific TaqMan primers (Roche Applied Science, Life technologies). The expression level was normalized to the expression of internal control gene GAPDH.

### Single-cell RNAseq analysis

BM Lineage-/Sca-1^+^/c-kit^+^ (LSK) cells from each experimental group of mice (pool of 3 mice per group) were FACS-sorted, diluted to 1,000 cells/μL, and 10,000 cells from littermate mice were loaded in each lane of a Chromium Controller Instrument (10x Genomics). Sequencing was performed on the Illumina HiSeq-4000 platform at a target depth of 230 million reads per simple. The resulting sequencing data from both 10x Genomics samples was aligned to genome version mm10 using Cell Ranger version 2.2.0 to obtain a filtered gene matrix files for further analysis. Approximately 5,000 cells were analyzed per group, with an average of reads per cell of approximately 50,000. These filtered matrices were supplied to AltAnalyze version 2.1.4 for joint unsupervised analysis with the software Iterative Cluster and Guide-Gene Selection (ICGS) version 2 (Ensembl version 72 database)^57^, using default parameters.6 ICGS2 identified 12 transcriptionally distinct cell populations, with all clusters containing generally similar proportions of cells from each capture. Cell-type annotations were obtained from ICGS2 biomarker and Gene Ontology gene-set enrichment predictions. UMAP visualization of hematopoietic stem and progenitor markers in AltAnalyze confirmed lineage assignments and suggested further heterogeneity of presumptive HSC-MPPs. To identify putative LT-HSC versus other HSC-MPP populations not in cell-cycle, we further subclustered these HSC-MPP (n=2,280 cells) using ICGS2 using a relaxed required marker gene cluster stability requirement (marker Pearson Cutoff = 0.20), to find four sub-clusters, denoted by distinct prior annotated HSC-subsets (gene-set enrichment of prior curated HSC marker genes). For both the sub-clustering and broader LSK clusters, we performed comprehensive cell-population differential comparison analysis with the software cellHarmony, comparing WT and AdipoQ-RXRα/β^Δ/Δ^ (empirical Bayes t-test p < 0.05, FDR corrected).

### Bulk RNAseq analysis

Around 10,000 SLAM HSCs (LSK/CD150^+^/CD48^−^) BM cells from each WT and AdipoQ-RXRαβ^Δ/Δ^ mouse were sorted in 100 µL of lysis buffer and RNA was isolated following manufacturer’s protocol (Arcturus Picopure RNA isolation kit, Applied Biosystems, # 12204-01). For bulk RNASeq analysis after ex vivo Resistin treatment, WT SLAMs were sorted and 1000 SLAMs/well were plated with either Vehicle or Resistin (10 ng/mL) in presence of SCF (100 ng/ml) and TPO (100 ng/mL) in X-VIVO 15 media. The plates were incubated for 16 hrs in 37 °C humidified CO_2_ incubator. After a brief centrifuge, the medium was carefully aspirated. Then the cells were lysed with 100 µL lysis buffer and the RNA was isolated following manufacturers protocol.

For Constitutive and Regulatory bone marrow adipocyte isolation, selected anatomical regions of tibia and femur were cut with a scalpel and bones were crunched. After adipocyte isolation, RNA was isolated using RNeasy kits, following manufacturer’s protocol (Qiagen, Germantown, MD; Cat #74104). Bulk RNAseq was performed using the adapted NEBNext Single Cell/Low Input RNA Library Prep Kit for Illumina (NEB, Ipswich, MA; Cat # E6420S). A range of 2 pg to 10 ng of purified input RNA was used for each sample. After the cDNA was amplified, quality and quantity of the amplified cDNA was measured using Agilent High Sensitivity DNA kit on a Bioanalyzer. 100pg-20ng of purified cDNA was used for the library preparation with TIGL’s own designed dual-index primers^58^.

After the sequencing library was made, it was sequenced using NextSeq P1-100 cycle (for ex vivo Resistin treated samples) or P2-100 (for in vivo WT and AdipoQ-RXRαβ^Δ/Δ^ mouse SLAM samples and bone marrow adipocyte samples), and cycle kit on the Illumina NextSeq 1000 sequencing system according to the manufacturer’s protocol with 50-nt read 1, 50-nt read 2, 8-nt index 1, and 8-nt index 2. The sequencing data analysis was performed with ALTAnalyze^57^.

A complete list of chemicals, antibodies and other reagents is presented in **Supplemental Table 5.**

## Statistical analysis

Data are presented as average ± standard deviation. Comparisons were performed with Student’s t test, chi-squared test and one-way or two-way ANOVA when required. GraphPad PRISM was used for statistical analysis. Statistical significance levels were established at 5%, 1% and 0.1%.

## Acknowledgements

This project has been funded by the National Institutes of Health grants R01 DK124115 and P01 HL158688 (JAC). The authors want to thank the Dana-Farber Cancer Institute Translational Immunogenomics Lab, Flow Cytometry Core Facility, Animal Facility and Confocal Microscopy Cores. The authors also want to thank the Cincinnati Children’s Hospital and Division of Experimental Hematology Animal Facility, Flow Cytometry and RNA sequencing cores.

## Supplementary Figure Legends

**Supplementary Figure 1:**
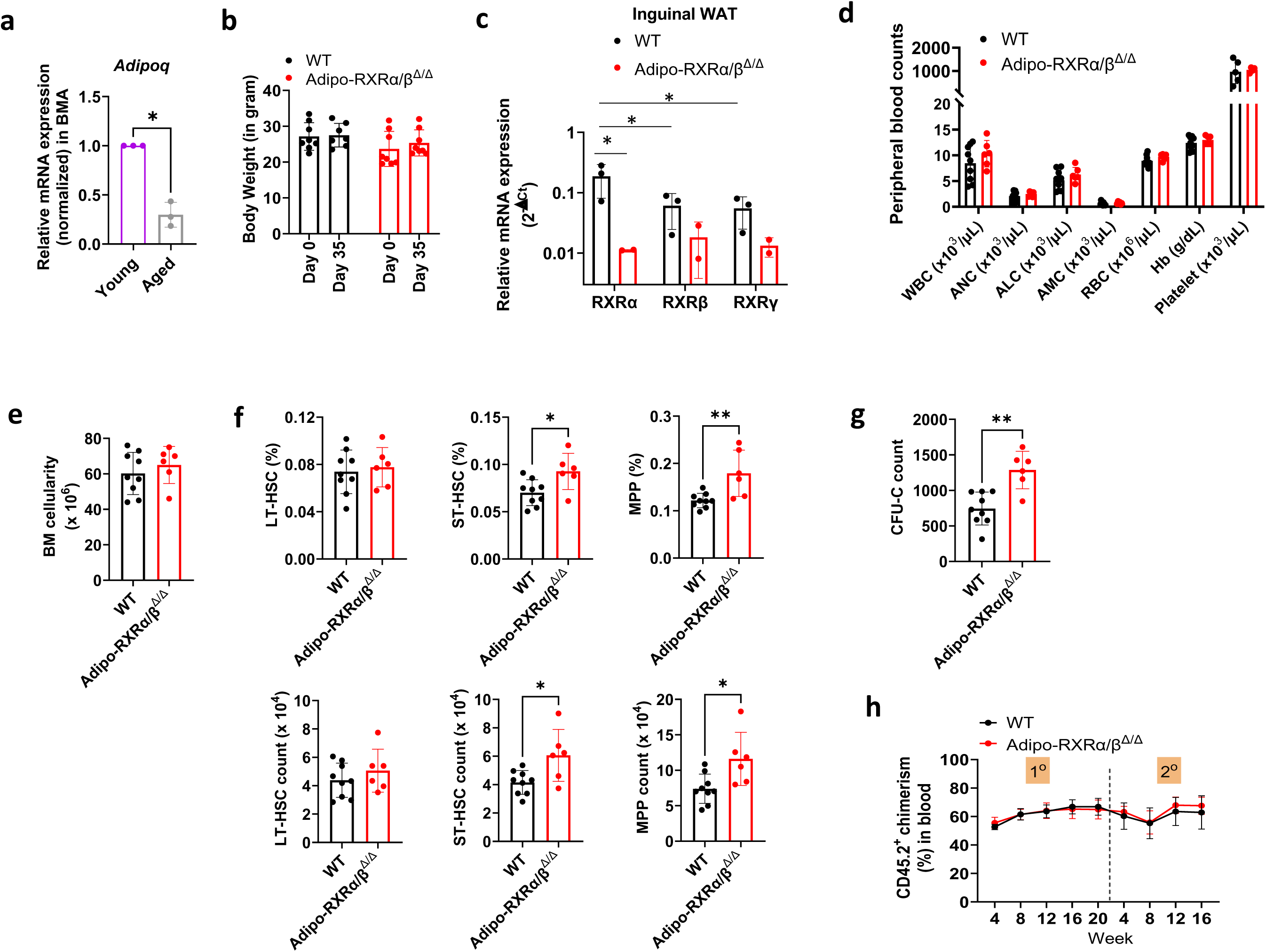
Related to Figure 1. (a) Relative mRNA expression of Adiponectin in BM adipocytes. (b) Body weight measurement before and after tamoxifen administration. (c) Relative mRNA expression of RXRs. (d) Blood parameters count in WT and Adipo-RXRαβ^Δ/Δ^ mice. (e) Quantification of total bone marrow cell in WT and Adipo-RXRαβ^Δ/Δ^ mice. (f) Quantification of LT-HSC, ST-HSC, and MPP percentages and numbers. (g) Quantification of CFU-C from bone marrow cells from WT and Adipo-RXRαβ^Δ/Δ^ mice. (h) Chimera analysis of primary and secondary transplanted mice. Data are presented as mean ± SD. Unpaired t-test and two-way ANOVA were performed for statistical analysis. *p<0.05; **p<0.01; ***p <0.001.

**Supplementary Figure 2:**
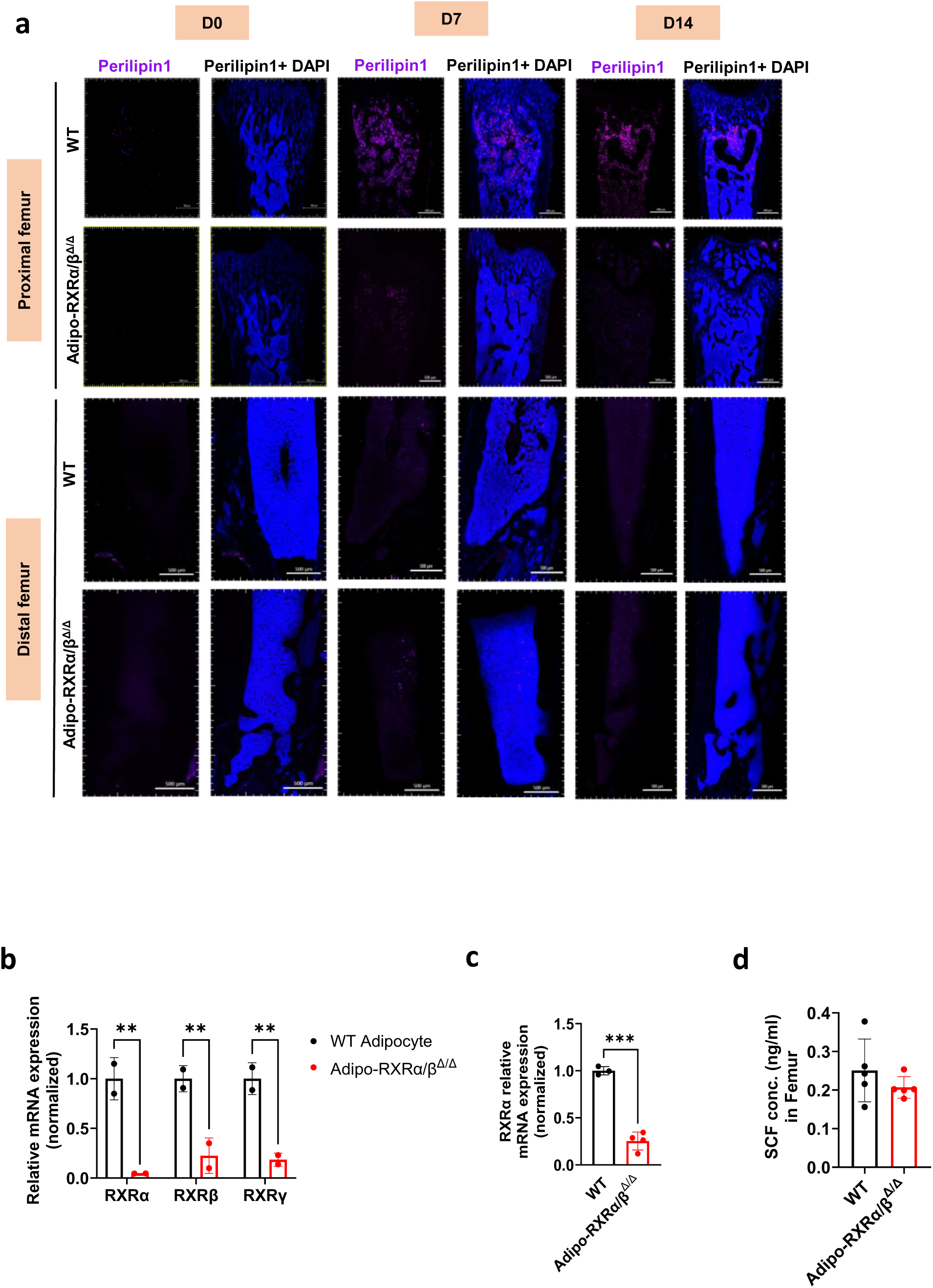
Related to Figures 1, 2 and 3. (a) Confocal images of Perilipin1 staining of the mouse femur after 5-FU administration. (b, c) Quantification of relative mRNA expression. Scale bar is 500 μm. (d) SCF concentration measurement of BM extracellular fluid by ELISA. Data are presented as mean ± SD. Unpaired t-test and two-way ANOVA were performed for statistical analysis.

**Supplementary Figure 3:**
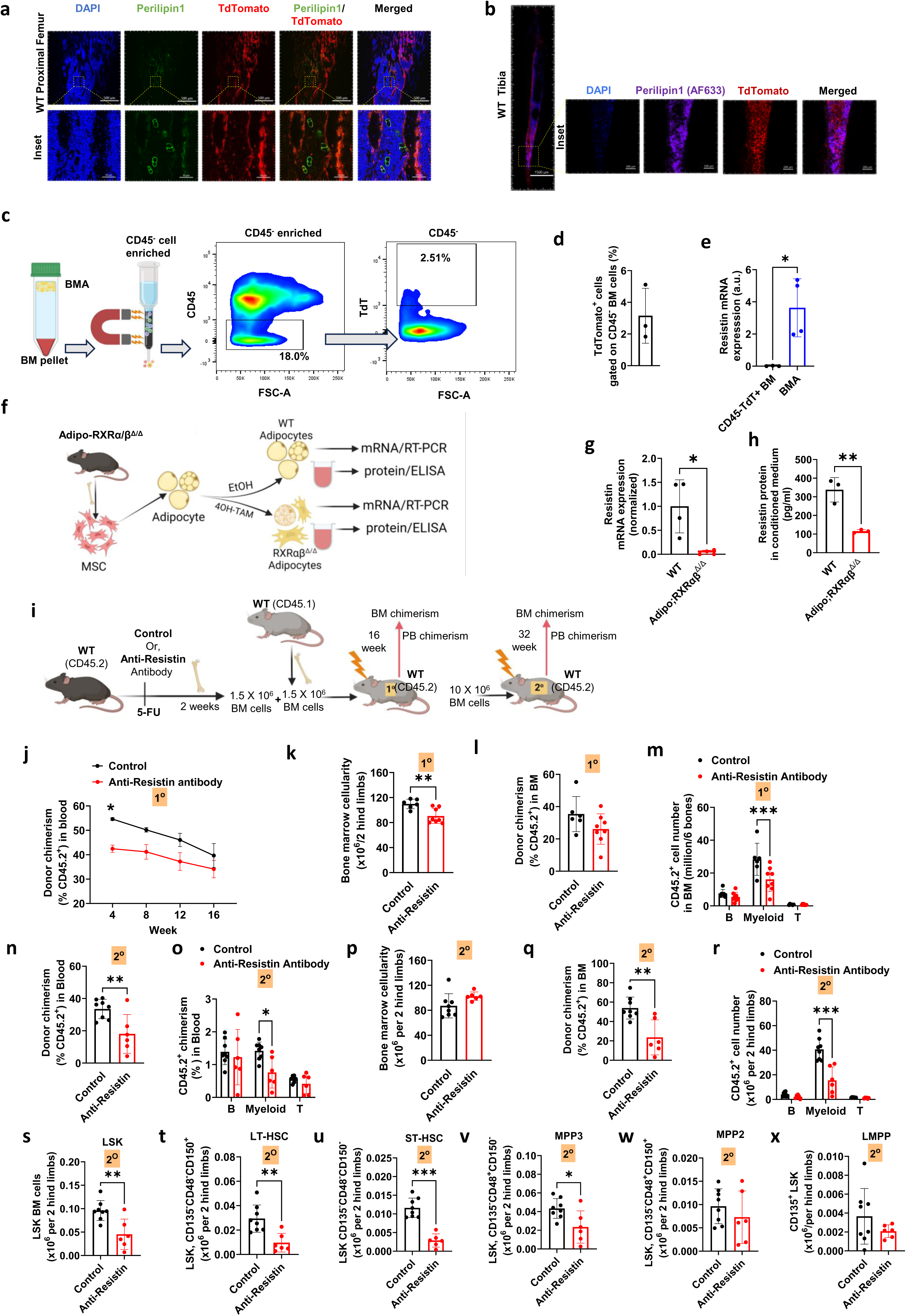
Related to Figure 3. (a) Confocal images of WT proximal BM TdTomato expressing adipocytes and preadipocytes in the proximal femur and (b) tibia. (c, d) Schematic representation of TdTomato+ (AdipoQ+) cell isolation with frequency. (e) Resistin mRNA quantification in the TdTomato+ (AdipoQ+) bone marrow stromal cells from WT mice and WT bone marrow adipocytes. (f) Schematic representation of MSC-derived adipocyte production ex vivo, RXRαβ^Δ/Δ^ adipocyte production, conditioned media, and cell mRNA isolation for Resistin quantification. (g) Resistin mRNA quantification. (h) Resistin concentration in the conditioned media. (i) Schematic representation of Resistin neutralization experiment. Quantification of (j) blood chimera, (k) bone marrow cellularity, (l) donor chimerism in bone marrow, (m) B, myeloid and T cell numbers in bone marrow in primary recipients. (n, o) Donor chimerism, B, myeloid, and T cell numbers in the blood of secondary recipients. Quantification of the (p) bone marrow cellularity, (q) donor chimerism in bone marrow, (r) B, myeloid, and T cell numbers in bone marrow in secondary recipients. (s-x) Quantification of different HSC/P populations. Data are presented as mean ± SD. Unpaired t-test and two-way ANOVA were performed for statistical analysis. *p<0.05; **p<0.01; ***p <0.001.

**Supplementary Figure 4:**
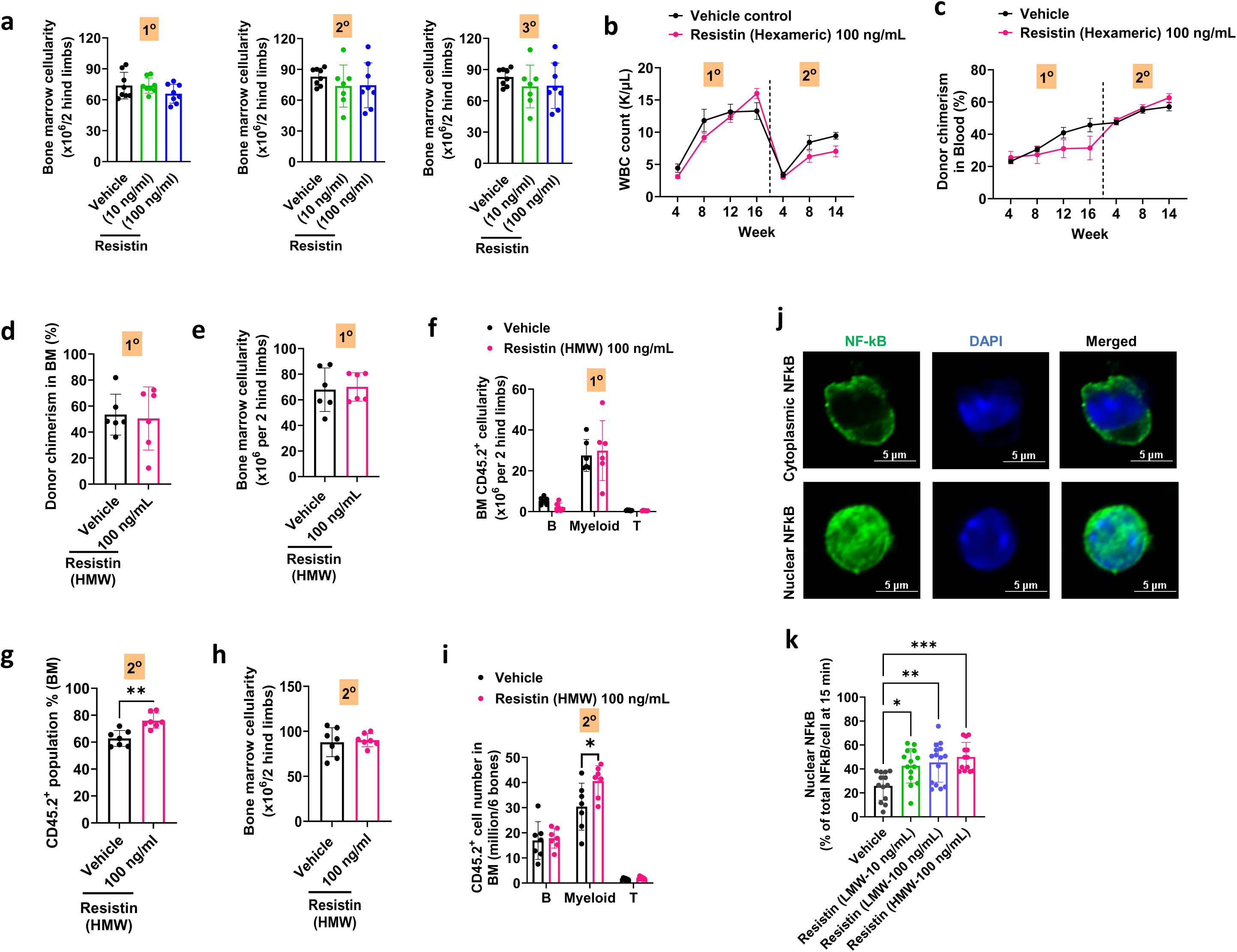
Related to Figures 4 and 5. (a) Bone marrow cellularity of primary and secondary recipients. (b) WBC (Leucocyte) count and (c) donor chimerism of primary and secondary recipients. (d-i) Donor chimerism, bone marrow cellularity, B, myeloid and T cell number of primary and secondary recipients. (j) Confocal microscopic images of HSCs with NF-kB staining. (k) Quantification of the NF-kB nuclear localization after Resistin treatment. Scale bar=5 μm. Data are presented as mean ± SD. Unpaired t-test and two-way ANOVA were performed for statistical analysis. *p<0.05; **p<0.01; ***p <0.001.

